# CESA: Cross-species Epitope Sequence Analysis for discovery of existing antibodies useful for phospho-specific protein detection in model species

**DOI:** 10.1101/2024.12.20.629730

**Authors:** Yanhui Hu, Chenxi Gao, William Mckenna, Baolong Xia, Majd Ariss, Stephanie Mohr, Norbert Perrimon

## Abstract

Signaling pathways play key roles in many important biological processes such as cell division, differentiation, and migration. Phosphorylation site-specific antibodies specifically target proteins phosphorylated on a given tyrosine, threonine, or serine residue. Use of phospho-specific antibodies facilitates analysis of signaling pathway regulation and activity. Given the usefulness of phospho-specific antibodies, a number of collections of these antibodies have been generated, typically for detection of phosphorylated mammalian proteins. Anecdotal evidence shows that some of these are also useful for detection of phosphorylated forms of orthologous proteins in model organisms. We propose that anti-phospho-mammalian protein antibody collections comprise an untapped resource for research in other species. To systematically analyze the potential utility of anti-phospho-mammalian protein antibodies in other species, we developed the Cross-species Epitope Sequence Analysis software tool (CESA). CESA identifies and aligns orthologous proteins in model species, then analyzes conservation of antibody target sites. We used CESA to predict what phospho-specific antibodies in a collection from Cell Signaling Technology (CST) might be useful for studies in *Drosophila melanogaster* and other species. CESA predicts that more than 232 sites on 116 *Drosophila* proteins can potentially be targeted by the antibodies initially developed at CST to detect human, mouse, or rat phosphoproteins.

## Introduction

Post-translational modification is essential for the regulation of many cellular processes. For example, phosphorylation can serve as a molecular switch for signal transduction by making the conformational change adjusting protein–protein binding surfaces or the localization of the target protein [1, 2]. The availability of phospho-specific antibodies has revolutionized the study of signal transduction process by making it possible to not only track the expression level but also the activity of the key players of signaling pathways. Phospho-specific antibodies are generated using peptides containing one or more phosphorylated amino acids and this process is more difficult than traditional antibody production which can be achieved by utilizing purified antigens or peptide immunogens, therefore, comparing to the available antibodies, the resources of phosphoantibodies is much limited. For example, based on antibodypedia [3], more than 7000 human genes can be targeted by antibodies while less than 500 human genes can be targeted by phospho-specific antibodies based on the PhosphoPlus database [4, 5], the largest phosphoantibody resource. The number dramatically drops for model organisms such as *Drosophila melanogaster, Danio rerio*, and *Caenorhabditis elegans*, for which only a handful phosphoantibodies are available commercially. To facilitate the identification of anti-phosphosite antibodies that might be useful in species other than the original target species, we developed the Cross-species Epitope Sequence Analysis (CESA) software tool to evaluate conservation of phosphorylated sites between the original species in which it was defined, e.g., human, and another species, such as a model organism species.

## Methods

### Tool development

The tool was developed in Python and DIOPT release 9 was used to map human, mouse, and rat genes to *Drosophila* genes using a filter to select only orthologous relationships with high and moderate ranks [6]. The tool communicates to the database using the SQLAlchemy package. The pandas package was used to process the query results. Protein global alignment was done using the pairwise2 package. The results were then filtered by selecting the longest isoform of each *Drosophila* ortholog gene (Table 2; Supplementary file 1). We also implemented a standalone version of the program for user to run the analysis locally without the need to connect to any database. This program is publicly available and can be run using a customized catalog file for species of choice.

### Information retrieval

The March 2024 release of the CST catalog file (Phosphorylation_site_dataset.gz) was obtained from https://www.phosphosite.org/staticDownloads.action. Ortholog mapping information along with NCBI Entrez Gene and RefSeq protein sequences were obtained from DIOPT release 9. Uniprot2geneid mapping was obtained from the UniProt knowledgebase (https://ftp.uniprot.org/pub/databases/uniprot/knowledgebase/idmapping/by_organism/). Human, mouse and rat protein sequences for UniProt records were obtained from UniProtKB (https://www.uniprot.org/uniprotkb).

### Antibody testing

5 × 10^6 S2R+ cells per well were plated into a 6-well plate in Schneider’s media supplemented with 10% FBS (fetal bovine serum) overnight. Media was aspirated and replaced with Schneider’s media containing 5 µg/mL insulin for 10 minutes. Cells pellets were lysed using lysis buffer containing protease and phosphatase inhibitors on ice for 10 minutes. Protein concentrations were measured by BCA protein assay (Thermo Fisher Scientific). 2-mercaptoethanol (final, 5%) and 4x Laemmli Sample Buffer were added to protein solutions and the mixtures were boiled for 10 minutes. Protein sample was loaded into each lane and the separation of proteins was performed in a 4–20% gradient Mini-PROTEAN TGX Stain-Free Precast Gel (Bio-Rad) using Novex™ Tris-Glycine SDS Running Buffer (Thermo Fisher Scientific). Proteins were transferred from gel to Immobilon® -FL PVDF Membrane (Millipore) with Pierce™ 10X Western Blot Transfer Buffer (Thermo Fisher Scientific) using a Trans-Blot® Turbo™ Transfer System (Bio-Rad) at 0.4A, 25V for 16 minutes. The membrane was blocked with 5% BSA in TBST for 1 hour at room temperature. The membrane was subsequently incubated overnight at 4C with Phospho-IGF-I Receptor β (Tyr1135/1136)/Insulin Receptor β (Tyr1150/1151) (19H7) Rabbit mAb #3024 (Cell Signaling Technologies) diluted 1:1000 in 5% BSA in TBST. The membrane was washed 3 times for 3 minutes with TBST before incubation with Goat anti-Rabbit IgG (H+L) Highly Cross Adsorbed Secondary Antibody, Alexa Fluor™ Plus 800 (Thermo Fisher Scientific) at a 1:2000 dilution and anti-Actin-Rhodamine at a dilution of 1:2500. The western blot was visualized using ChemiDoc MP Imaging System (Bio-Rad).

### Code availability

The standalone version of CESA program is available at GitHub (https://github.com/chenxi-gao/antibody_discovery).

## Results

### Defining rules for predicting cross-reactivity of phospho-specific antibodies with orthologous proteins in non-target species

We reasoned that for an antibody to be useful to detect a corresponding phosphosite on an orthologous protein in another species, the amino acid sequence of the phosphosite and the surrounding region must be conserved. However, examples in the literature [7] and as reported in commercial antibody catalogs (e.g., selected examples in Table 1) suggest that the region of conservation surrounding the phosphosite can be as small as 6 amino acids.

**Table 1.**
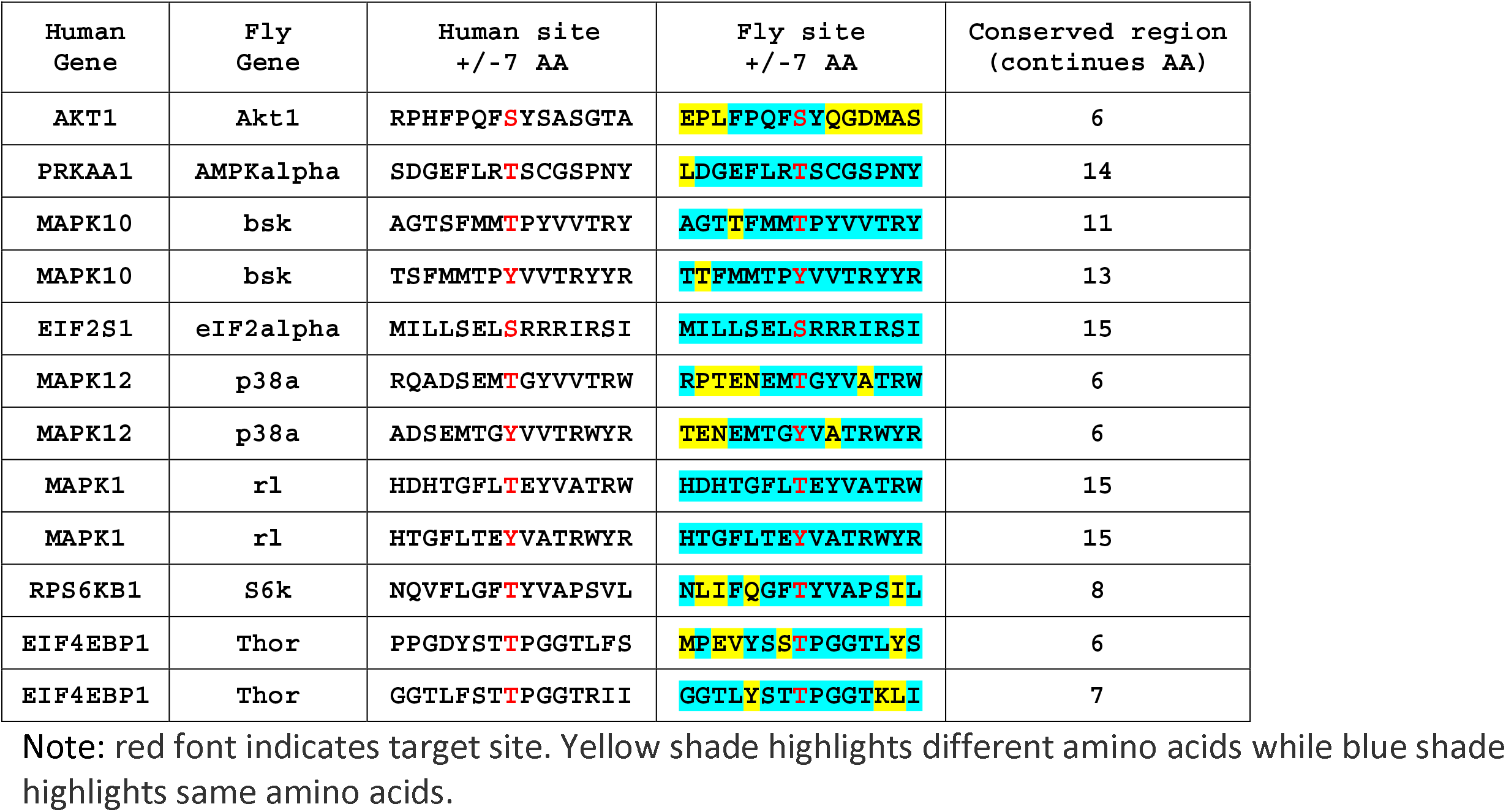
Commercial antibodies against mammalian proteins with validated reactivity in Drosophila.

To more precisely infer what amino acid identity and similarity is sufficient to predict cross-reaction of an anti-phosphosite antibody directed against ‘protein A’ with some orthologous ‘protein B’ in another species, we first analyzed attributes of anti-phosphosite antibodies already reported to cross-react with another species. As a source for information, we turned to the PhosphoSitePlus web-based bioinformatics resource (https://www.phosphosite.org) [4, 5], which is made available by Cell Signaling Technologies (CST), one of the commercial sources for anti-phosphosite antibodies. The March 2024 PhosphoSitePlus release file lists 378,971 phosphosites annotated in 27 organisms, 2357 of which are reported to be recognized by CST antibodies. Among the 2357 antibodies, 12 antibodies are reported in PhosphoSite Plus to cross-react with phosphosites on 8 *Drosophila* proteins (Table 1). There are a few other commercial sources with a limited number of anti-phosphosite antibodies that can cross-react with *Drosophila* as well, however, the information regarding the target site is quite limited, for example, the protein accession as well as the peptide sequence information is not public available. Therefore, we only focused on the antibodies from CST. After we aligned the original target sites with 7 amino acids on each side with the protein of *Drosophila* ortholog, we evaluated the conservation for these 12 validated sites of cross-reactivity. As expected, the aligned sites are conserved between human and *Drosophila* orthologs, with identical sequences over 6-15 amino acid region spanning the phosphosite region.

### Development of the Cross-species Epitope Sequence Analysis (CESA) software tool

We developed the Cross-species Epitope Sequence Analysis (CESA) software tool to evaluate conservation of phosphorylated sites between the original species in which it was defined, e.g., human, and another species, such as a model organism species, with the ultimate goal of identifying anti-phosphosite antibodies that might be useful in species other than the original target species. There are five major steps in the pipeline (Figure. 1A). First, CESA maps the epitope sequence to a current protein release so that the full protein sequence can be identified for alignment and the amino acid position of the target site relative to the reference protein sequence is noted. This step of validating the information about the target site on source protein, against which the antibody was originally generated, is critical for the downstream analysis, however, we observed that some of the protein accession numbers in the CST catalog are outdated, therefore failed the step retrieving full protein sequence from current protein resources. To overcome this issue, we added an additional process for records failed the first step, to identify the current protein records from RefSeq database using gene symbol provided in the catalog as well as the gene2refseq mapping file from NCBI (link). Then compare these protein sequences retrieved based on gene symbol with the phospho peptide sequence from the catalog to annotate the location of the phosphosite (Figure. 1B). Second CESA maps gene symbols and/or protein accession numbers to NCBI Entrez Gene identifiers using the gene identifier mapping tools developed in house. This allowed us to assign Entrez Gene IDs, which are needed to query orthologous genes using DIOPT, an ortholog mapping database that integrates ortholog mapping from more than twenty algorithms [6]. Third, CESA maps input genes from such as human or mouse to orthologs in the second species such as *Drosophila* based on ortholog predictions from DIOPT with the filter selecting only the mapping of high or moderate confidence [6]. Next, CESA performs pairwise protein alignment of each input protein sequence with all isoforms of the model species ortholog using a global alignment algorithm. At the last step, a parser was developed to identify the target sequence of the input protein and the aligned *Drosophila* sequence, then analyze the conservation of all peptides 6 to 11 amino acids long spanning each amino acid that can be phosphorylated to identify the sites that are identical between the two species. We allow only one conserved amino acid replacement at the location not right next to the target amino acid (S/T/Y). The amino acid conservation in the area immediately surrounding and including the aligned phosphorylation site with different peptide length cutoff is summarized in the output file.

**Figure 1.**
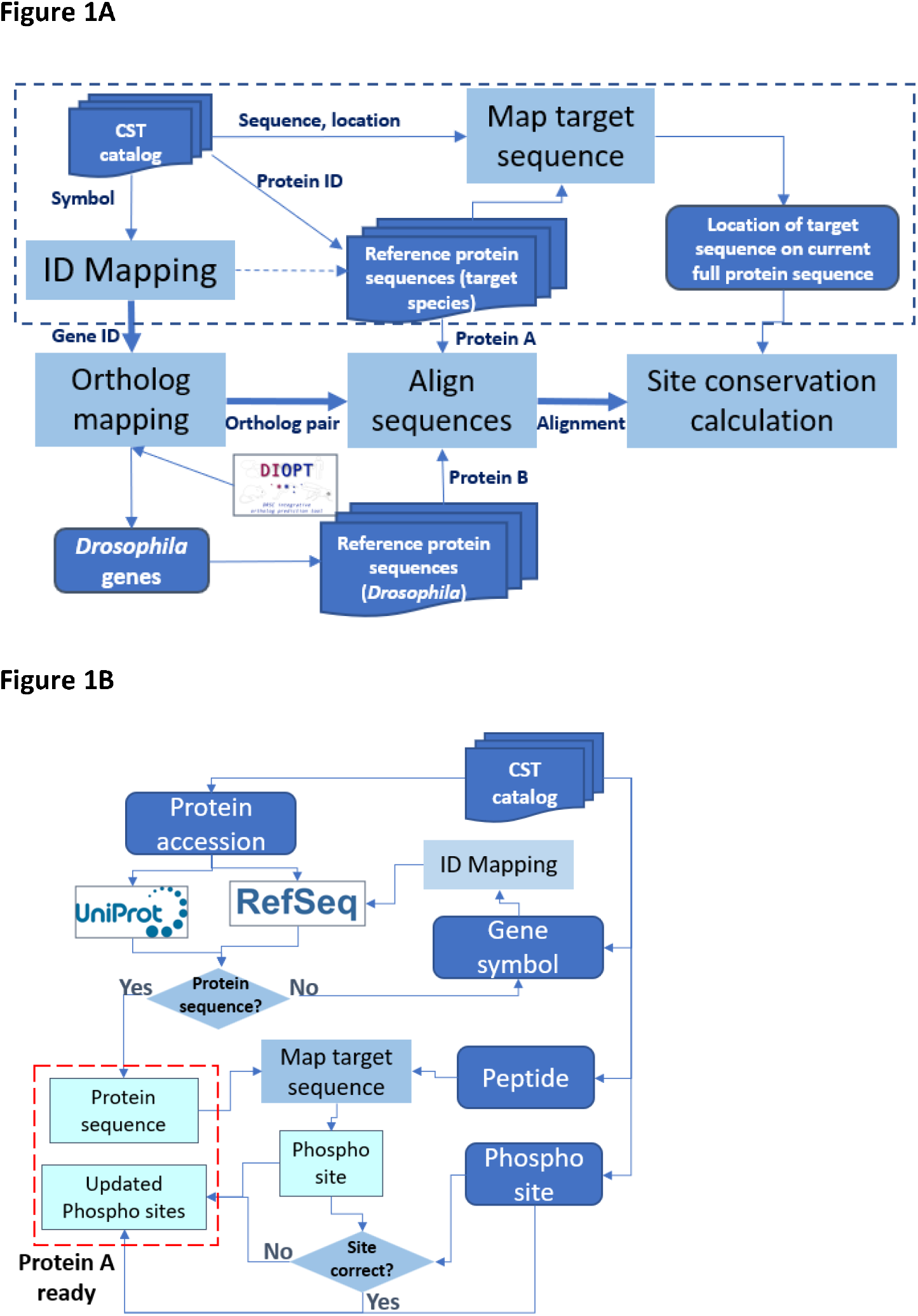
Design of the Cross-species Epitope Sequence Analysis (CESA) software tool. **A**. Overall workflow. Protein A, original target. Protein B, putative orthologous target identified in another species. An abbreviated presentation of steps needed to get some ‘protein A’ ready for analysis is shown within the dotted line box. Dark blue represents for input and output lists while light blue represents major steps. **B**. Detailed presentation of processing steps needed to get some ‘protein A’ ready for analysis.

### Application of CESA to predict Drosophila targets of phosphosite antibodies

As a test case, we applied the CESA pipeline to identify anti-phosphosite antibodies at CST that might cross-react with an orthologous protein in *Drosophila*. As an input set, we selected the subset of 2,301 phosphorylated sites in PhosphoSitePlus that were originally annotated for human, mouse, or rat and can be targeted by antibodies in the CST collection. Using CESA, 998 of 1,170 genes from human, mouse, and rat were mapped to 659 *Drosophila* genes, among which, 232 phosphorylation sites on 116 *Drosophila* genes that can be potentially targeted by the CST antibodies originally developed for other species using 6 amino acids as cut-off (Table 3, sup table 1). Among the 116 *Drosophila* genes that can be potential targeted by CST antibodies, there are only 3 unannotated genes with CG number as gene symbol, and 87 (75%) of them have more than 20 associated publications (sup table 1). Gene set enrichment analysis was done using PANGEA [8] to identify the gene groups that are over-represented among them and the top gene groups are kinases (47 genes, P adj = 2.80E-24), core components of major signaling pathways (29 genes, P adj = 1.21E-20) (Figure 2, sup table 2). For example, *Drosophila* insulin receptor (InR, FBgn0283499) is one of the core components of Insulin signaling pathway. The region spanning the three C-terminal tyrosines (Y1545/1549/1550 on the RB isoform) of this gene, not only shares 100% identity with human, but also the phosphorylation at these sites has been observed in *Drosophila* based on the phospho-proteomics data in iProteinDB [9] (Sup Figure 1a). The Abcam antibody ab62321 has been tested in human for the 1^st^ site while the CST antibody CST3024 has been tested in human, mouse and rat for the 2^nd^ and 3^rd^ site. Based on literature, Abcam antibody ab62321 has been tested and shown reactivity in *Drosophila* S2R+ cells upon the insulin treatment, which activates Insulin signaling pathway [7]. Therefore, very likely the CST antibody CST3024 can also target the phosphorylated *Drosophila* protein. Indeed, using the same protocol of S2R+ cell handling, we tested antibody CST3024 and the result shows reactivity as well (Sup Figure 1). It is reasonable to suspect that the antibodies identified by CESA will be important reagents for *Drosophila* research (Table 1).

**Figure 2.**
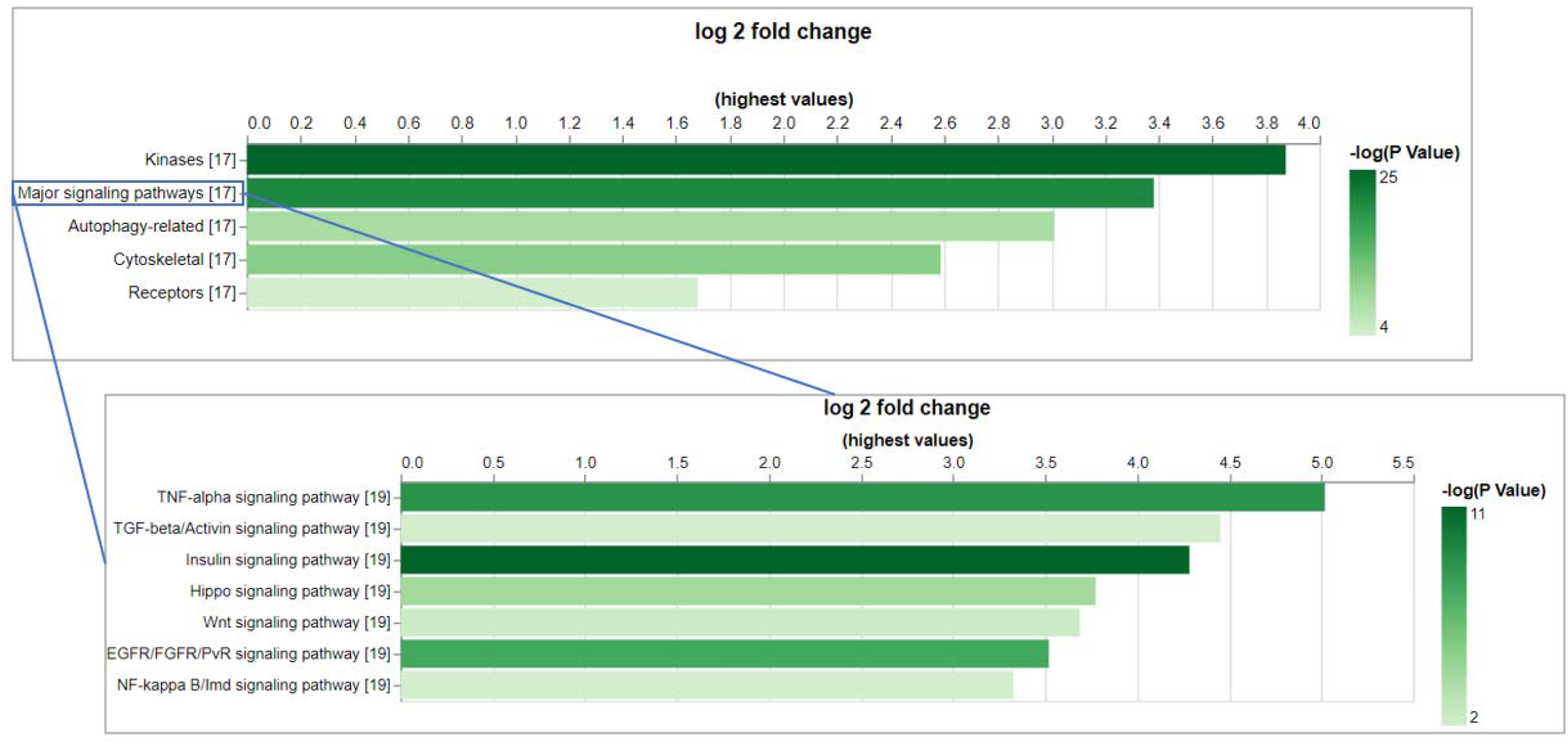
The gene set enrichment analysis (GSEA) of the *Drosophila* genes that can be potentially targeted by CST phospho-antibodies indicates the over-representation of core components of major signaling pathways. GSEA was done using PANGEA and the gene sets annotated by GLAD as well as PathOn (signaling pathway core components) were selected. Enrichment result was filtered using P value 0.05 as cutoff and on the bar graphs from PANGEA, the height reflects fold enrichment while the darkness reflects the negative log10 pvalue of enrichment.

In addition, we also ran CESA for Zebrafish (*D. rerio*), frog (*X. tropicalis*), mosquito (*A. gambiae)*, worm (*C. elegans*) and we identified between 584 to 75 genes, respectively, that can be potentially targeted by the current CST collection (Table 3, sup table 2). We observed that a larger portion of phospho antibodies work cross species between genetically closer species, such as 57% of human phosphosites conserved with Zebrafish, compared to only 17% of human phosphosites conserved with *Drosophila* (Table 2). Nevertheless, CESA systematically identifies the potential reagents from existing collection, which is significantly more than the reagents currently available commercially. Altogether, our analysis should advance research in relevant communities, particularly for species with very limited antibody resources.

**Table 2.** Summary of phosphosites reported in PhosphoSitePlus, availability of corresponding CST antibodies and their conservation with *Drosophila, zebrafish, frog, mosquito* and *c. elegans*.

| Species | Sites | Genes | Sites with antibody | Genes with antibody | Sites conserved in fly | Sites conserved in frog | Sites conserved in fish | Sites conserved in mosquito | Sites conserved in worm |
| --- | --- | --- | --- | --- | --- | --- | --- | --- | --- |
| human | 240,118 | 17,467 | 1,019 | 485 | 187 | 658 | 599 | 181 | 114 |
| mouse | 105,223 | 13,475 | 774 | 407 | 171 | 514 | 485 | 160 | 107 |
| rat | 31,721 | 7,261 | 508 | 278 | 116 | 328 | 311 | 111 | 72 |
| other | 1,893 | 676 | 56 | 33 | NA | NA | NA | NA | NA |

**Table 3.** Known or predicted phosphosites in *Drosophila*, zebrafish, frog, mosquito and *C. elegans* that can potentially be targeted using CST antibodies developed to target phosphosites in mammals.

| Cut-off of conserved region (aa) | Fly<br>*D. melanogaster* | Frog<br>*X. tropicalis* | Zebrafish<br>*D. rerio* | Mosquito<br>*A. gambiae* | Worm<br>*C. elegans* |
| --- | --- | --- | --- | --- | --- |
| 6 | 115 | 472 | 599 | 97 | 76 |
| 7 | 111 | 458 | 578 | 91 | 73 |
| 8 | 105 | 446 | 564 | 87 | 70 |
| 9 | 97 | 431 | 544 | 83 | 68 |
| 10 | 92 | 418 | 518 | 77 | 67 |
| 11 | 88 | 399 | 495 | 73 | 67 |

## Discussion

Antibodies targeting the phosphorylated peptide of endogenous protein are crucial to study the signaling pathways for processes like cell division and migration but developing such reagent is both labor and time consuming. Such resource is quite limited for majority of model organisms. While current commercial collections were originally designed for mammalian proteins, they may also be useful for detecting phosphorylated proteins in other organisms. To explore this potential, we developed CESA tool to help researcher to identify such reagents. Although we mainly used *Drosophila* as an example in this study, the CESA approach can be applied to any other model and non-model organisms. It is worth noting that since CESA considers all isoforms in the second species, it can predict multiple bands on Western blots if conserved sites are identified in more than one protein isoform of different lengths. It is also possible to configure CESA to predict cross-gene activity based on paralog relationships. In addition, CESA can be adapted for other applications, such as analyzing the epitope sequences of regular (non-phospho specific) antibodies. In summary, we systematically evaluated current CST phospho antibody to predict the cross-species activity in Drosophila and a few other organisms, and identified significant number of new potential useful reagents for species with very limited antibody resources. The result will likely advance research in relevant communities.

## Acknowledgements

We appreciate the feedback from the Perrimon lab for their valuable feedback. We extend our gratitude to the Harvard Medical School Research Computing and IT-Client Services teams for their consultation, web hosting, and support. The CST antibodies were generously provided by Roberto Polakiewicz and Sean Beausoleil. This article is governed by HHMI’s Open Access to Publications policy. HHMI lab heads have previously granted a nonexclusive CC BY 4.0 license to the public and a sublicensable license to HHMI for their research articles. Accordingly, the author-accepted manuscript of this article will be made freely available under a CC BY 4.0 license immediately upon publication.

## Author Contributions

Conceptualization, N.P. and M.A.; methodology, Y.H.; software, C.G.; validation, W.M. and B.X.; formal analysis, Y.H.; resources, M.A.; writing—original draft preparation, Y.H.; writing— review and editing, S.M.; visualization, Y.H.; supervision, N.P.; funding acquisition, N.P. All authors have read and agreed to the published version of the manuscript

---

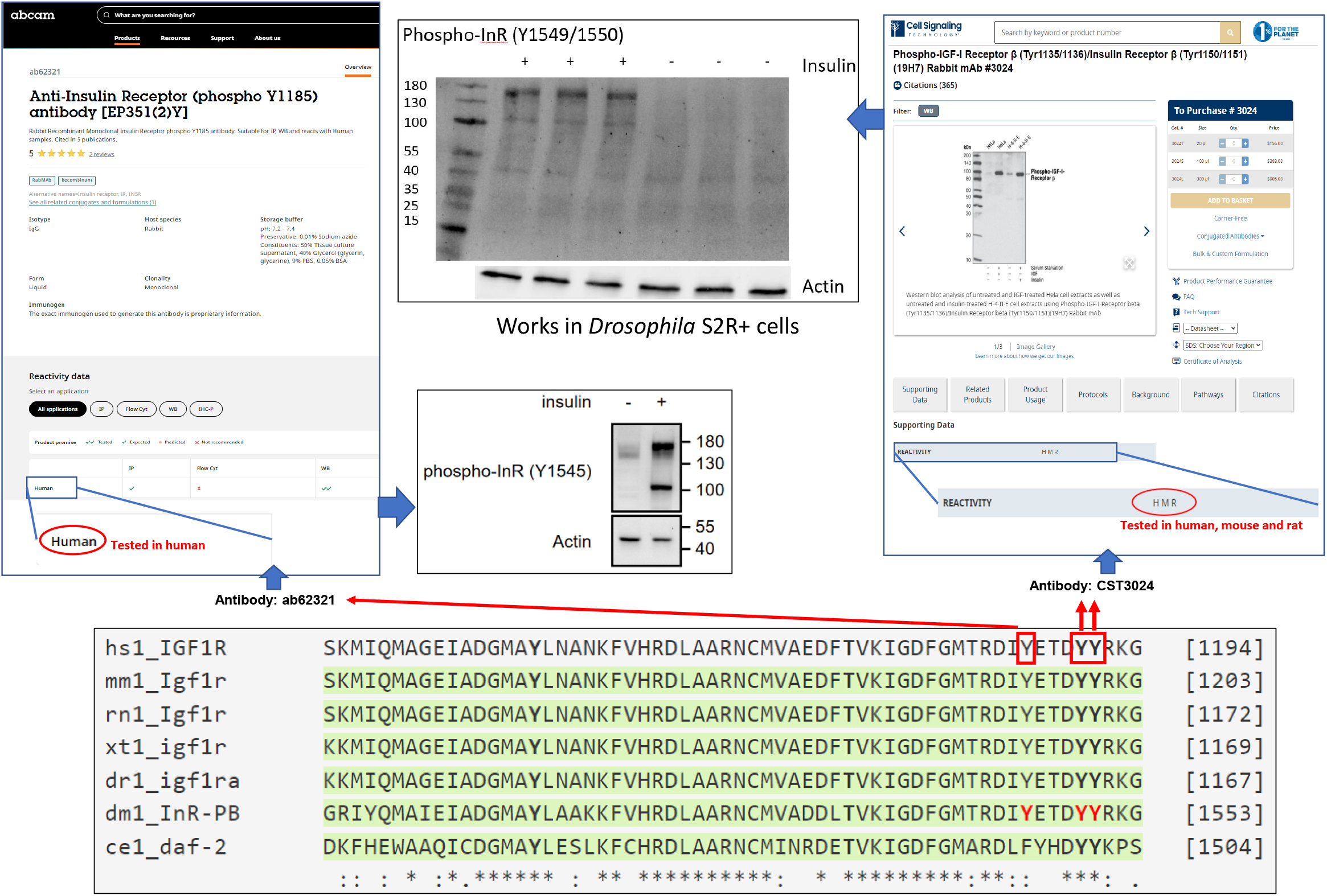

